# *Centris pallida* (Hymenoptera: Apidae) male body size decreases across five decades

**DOI:** 10.1101/2022.05.27.493633

**Authors:** Meghan Barrett, Meredith G. Johnson

**Affiliations:** Department of Biology, California State University Dominguez Hills, Carson, CA 90747; School of Life Sciences, Arizona State University, Tempe, AZ 85281

## Abstract

Historical data suggest that many bee species have declined in body size. Larger-bodied bees with narrow phenological and dietary breadth are most prone to declines in body size over time. This may be especially true in solitary, desert-adapted species that are vulnerable to climate change – such as *Centris pallida* (Hymenoptera: Apidae). In addition, body size changes in species with size-linked behaviors could threaten the prevalence of certain behavioral phenotypes long-term. *C. pallida* solitary bees are found in the Sonoran Desert. Males use alternative reproductive tactics (ARTs) and are dimorphic in both morphology and behavior. *C. pallida* male body size has been studied since the 1970s in the same population. We collected body size data in 2022 and combined it with published records from 1974-2022. We find a persistent decline in the mean head width of patrolling males, and shifts towards smaller body sizes in the populations of males found foraging and hovering. Both morphs declined in average body size, and the proportion of large-morph males in the population decreased by 8%. Mating males did not decline in mean body size over the last five decades. We discuss hypotheses related to the decline in *C. pallida* male head width. Finally, we advocate for *C. pallida* as an excellent study system for understanding the stability of ARTs with size-linked behavioral phenotypes.

## Introduction

Anthropogenic changes in climate and habitat have caused a decline in abundance, a shift in geographic range or phenology, and changes in the morphology, physiology, or behavior of many species, sometimes leading to alterations in life history and ecological relationships (Bartomeus et al. 2011, Biesmeijer et al. 2006, Jacobson et al. 2018, Béltran et al. 2021, Burraco et al. 2020, Chou et al. 2019, Chung & Schulte 2020, Duffy et al. 2015, Huey & Kingsolver 2019, Kuhlmann et al. 2012, Ockendon et al. 2014, Parmesan 2006, Sinervo et al. 2010, Walters & Hassall 2006, Zattara & Aizen 2021, Sánchez-Bayo & Wyckhuys 2019, Burkle et al. 2013, Turley et al. 2022).

Bees may be particularly vulnerable to climate change or habitat modifications. As heterothermic ectotherms they rely on environmental conditions to maintain non-lethal body temperatures (Wieser 1973, Huey & Stevenson 1979). This is compounded by a reliance on resources (e.g., nesting resources, plants) that are often disturbed by human activity (Grab et al. 2019, McCabe et al. 2021). Documented changes in bee populations over time can help determine how human activity may impact bees with different life history characteristics, with knock-on effects for their ecological relationships. (e.g., Cane et al. 2006, Burkle et al. 2013). For example, studies in Europe and the Northeastern regions of North America have demonstrated that larger-bodied bee species are more likely to experience declines in population (Bartomeus et al. 2013, Scheper et al. 2014, Oliveira et al. 2016), or stronger intraspecific declines in body size (Nooten & Rehan 2020, but see Gérard et al. 2019), perhaps due to greater nutritional requirements (Müller et al. 2006).

Bee body sizes have not changed uniformly over time across species, sexes, or regions (Nooten & Rehan 2020, Bartomeus et al. 2013, Scheper et al. 2014, Oliveira et al. 2016, Kleijn & Raemakers 2008), suggesting life history traits and ecosystem characteristics (e.g., habitat fragmentation, Warzecha et al. 2016) may play important roles in determining the impact of human activities on body size. For most bees, body size is determined by resource provisioning in the larval stage (Chole et al. 2019, Kukuk 1996, Alcock 1984, Lawson et al. 2017), and can thus be strongly influenced by changes in resource availability (Chown & Gaston 2010). Bees with narrow phenological windows and bees that are dietary specialists are likely to be more susceptible to declines, given their reliance on a smaller pool of available resources (Bartomeus et al. 2013). Solitary and desert-adapted bees are expected to be particularly vulnerable to climate change (Sala et al. 2000, Loarie et al. 2009, Vale & Brito 2015, Hamblin et al. 2017, Burdine & McCluney 2019, McCabe et al. 2021, but see Silva et al. 2018), given the thermal and hygric stressors already evident in their environment. To date, no studies have tracked changes in morphology of any solitary bees in desert ecosystems. Declines in body size could also generate species-level behavioral changes whenever size contributes to behavior, such as in many alternative reproductive tactic systems (ARTs). ARTs occur when categorical variation in the morphological/behavioral traits is associated with mating across individuals of the same sex within the same population (Oliveira et al. 2008, Paxton 2005, Shuster 2010). In many such systems, larger- and smaller-bodied individuals use different strategies for accessing mates (e.g., fighters vs. sneakers; Oliveira et al. 2008). Declines in body size may alter the occurrence of size-linked morphs, and their behaviors, resulting in a loss of important intraspecific variation over time.

*Centris pallida* (Hymenoptera: Apidae) are widespread, common solitary digger bees are found in the Sonoran Desert of the Southwestern United States and Northern Mexico. This species often forms dense nesting aggregations, where many thousands of individuals occupy several hectares. *C. pallida* mating aggregations may persist in the same locations over multiple decades – allowing for the continuous resampling of a population (Barrett 2022). Males emerge first from their natal nests at these sites and use ARTs to find mates. Males are behaviorally and morphologically dimorphic, using different sensory mate location strategies and microclimates (Alcock 1976, Alcock et al. 1977a, Snelling 1984, Barrett et al. 2021, Barrett 2022). Large-morph, ‘metandric’ males with pale gray coloration patrol in sinuous loops ∼10 cm over the emergence site and use scent to locate females emerging from natal nests; males then engage in fights for the opportunity to dig up and mate with emerging females (Alcock et al. 1976b). Small-morph males, with dark brown coloration, are more behaviorally flexible; they may patrol, but are often found hovering near plants where they use visual cues to locate females or mating pairs flying away from the aggregation site that may be interrupted. Large-morph males have a clear fitness advantage in situations where mating aggregations are densely populated (Alcock et al. 1977a, 1977b, Alcock 1984, Alcock 1995, Alcock 2013a), as is always the case in this specific *C. pallida* population.

*C. pallida* are large-bodied, desert-adapted, solitary bees with an extremely narrow phenological and dietary breadth. Bees nest for less than six weeks in the late spring, and females utilize four species of flowering trees for larval nectar/pollen provisions (*Parkinsonia microphylla, P. aculeata, Olneya tesota*, and, *rarely, Psorothamnus (Dalea) spinosa;* Alcock et al. 1976a). *P. microphylla* is the predominant host plant for *C. pallida* over its entire geographic range (Buchmann, pers. comm). The body sizes of male *C. pallida* have been studied at the same site since for 48 years (Alcock et al. 1977a). By combining historical data on *C. pallida* body sizes from 1974 – 2018 (Alcock et al. 1977a, Alcock 1984, Alcock 1989, Alcock 2013a, Barrett 2022) with data we collected in 2022 from the same population, we aimed to determine if male *C. pallida* bees have experienced persistent changes or persistent stability in body size. Our results demonstrate that the population of males found foraging at trees (representative of the total population of males), hovering, and patrolling have all declined in body size since 1974. However, the population of mating males has not persistently declined in body size over time. In addition, we present the first body size data on female *C. pallida* from this site, for use in future historical comparisons. Finally, we discuss several hypotheses of causes for the decline in male body size, and the implications of declining male body size on the stability of the *C. pallida* ART.

## Methods

### Head width measurements

We collected head widths at the same site where most previous research on *C. pallida* body size has been conducted (Alcock et al. 1977a, Alcock 1984, Alcock 1989, Alcock 2013a), in the floodplain by Blue Point Bridge/Saguaro Lake over the Salt River north of Mesa, Arizona (33.552, -111.566). Alcock et al. (1977a) demonstrated that head width is correlated with body weight in *C. pallida* males, and serves as a reliable field indicator of body size.

We collected bees (n = 921 males, 114 females) in the same manner as described in Alcock (1984, 2013), to be sure our data were comparable. Briefly: for patrolling males, we made rapid sweeps low to the ground with an insect net through open areas of searching males. We approached and collected digging/fighting males by hand. We collected foraging males and females with a telescoping insect net as they visited paloverde trees (*Parkinsonia* sp.). We collected mating males and females while engaged in copulation on the ground or vegetation. We collected hovering males near mesquite or palo verde trees, wherever they had established stable aerial stations. We measured head widths to 0.01 mm using digital calipers (Wen 10761), before releasing the bees. We collected bees daily when the aggregation was most active between April 20 and May 8, 2022, generally between the hours of 0700 and 1130, and noted the time of collection and behavior for each individual.

### Historical data for comparisons

To obtain historical male head width data, we used reported means and standard deviations in Alcock et al. 1977a, Alcock 1984, Alcock 2013a, and Barrett 2022. Together, these studies report data collected in: 1974, 1975, 1976, 1982, 1988, 2011, 2012, and 2018. As prior surveys of the population by Alcock (et al. 1977; Alcock 1984, 1989, 2013a) generally only captured bees between roughly 0700 and 0930, and did not note the specific individual time of capture. We therefore also used only those bees captured before 0930 in our sampling (2022) for the historical comparison with prior literature. Barrett (2022) reported data for mating, hovering, and patrolling bees from this site in 2018, but patrolling bees were not captured in a manner consistent with Alcock’s prior surveys and so we elected to only use the mating and hovering male datasets in our current comparison.

### Changes in the Proportion and Size of Each Morph

To determine changes in the proportion of large- and small-morph males in the total population (represented by males caught foraging, Alcock et al. 1977a), we used the head width frequency distributions. These distributions are bimodal, with the two local maxima representing the two morphs; there is thus a local minimum in frequency between the small- and large-morph males (see Fig 2, where an arrow indicates the local minimum). The proportion of all males in the foraging population that were smaller in head width than the local minimum was assumed to be the proportion of small-morph males in the population. Those males larger in head width than the local minimum were assumed to be large-morph males. The proportion within the local minimum were considered the rare intermediate males that occur in all *Centris* species with male dimorphism (only 5-6% in both the 1974/5 and 2022 samples).

To determine morph-specific changes in body size, we assessed shifts in the head width frequency distributions on either side of the local minimum. We corroborated this shift in small-morph male body size by looking at changes in the head widths of hovering males, specifically. Hovering is completed *only* by small-morph males, and therefore can be used as a secondary method for assessing body size changes within this morph.

### Statistical analysis

Data were analyzed using GraphPad Prism 9.3.1 (GraphPad Prism for Windows 2021). When analyzing data on the total population in 2022 (as compared to the before-0930 dataset used for the historical data comparison), we include all bees captured in 2022 (including males for which we had not taken data on time of capture). Because head width data were not normally distributed for all ehavior categories, we used a Kruskal-Wallis test with post-hoc Dunn’s MCT to analyze differences between hovering, patrolling, and mating male populations. We used a Mann-Whitney test to compare mating and foraging male populations, and an unpaired t-test to compare digging and fighting males and mean hovering male head widths from 1974/1975 and 2022. We used a linear regression to analyze the effect of time of day on patrolling and hovering male head widths. We also used a linear regression to analyze the effect of year on mean patrolling and mating male head width across studies, using the reported mean, standard deviation (SD), and population size (n) of each study. All data can be found archived on Dryad (Barrett & Johnson 2022).

## Results

### *C. pallida* head widths in 2022

The ranges and frequencies of male head widths for the 2022 population of males engaged in foraging, patrolling, hovering, mating, digging, and fighting behaviors are shown in Figure 1 (mean ± SD and n, reported in Supplementary Table 1). We also present mean female head widths in this population, for females caught while mating, foraging, or nesting in Supplementary Table 1 (distribution of female head widths: Supplemental Figure 1).

**Figure 1.**
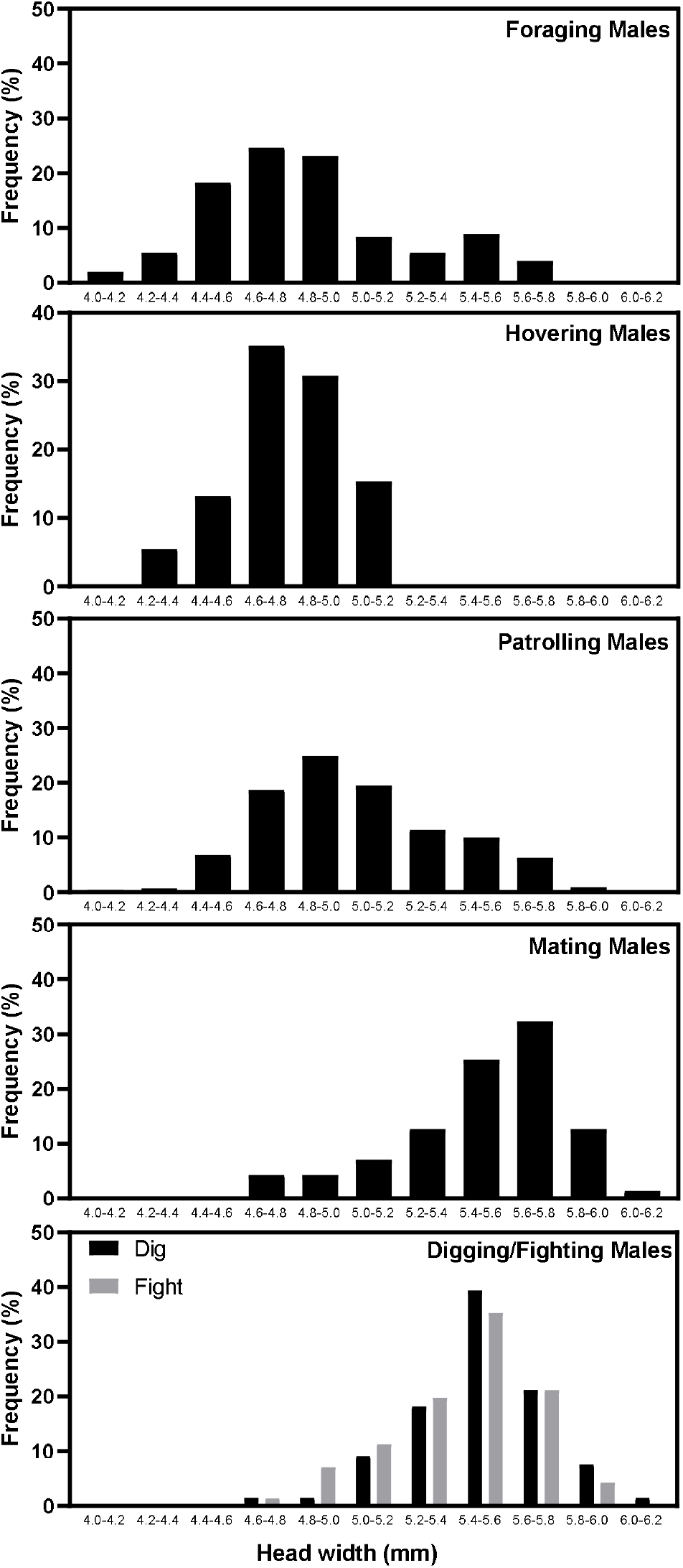
Distribution of male head widths by behavioral category in 2022. Distribution of male head widths (n = 921) collected while foraging (n = 203), hovering (n = 91), patrolling (n = 410), mating (n = 71), and digging/fighting (n = 66/71) before 1130 on April 20 – May 8, 2022 at Blue Point Bridge, Arizona.

Hovering males were significantly smaller than patrolling males (Figure 1; Kruskal-Wallis: K-W = 140.4, p < 0.0001; Dunn’s MCT: Z = 6.51, p < 0.0001), and both types of males were smaller than mating males (patrolling vs. mating: Z = 8.70, p < 0.0001; hovering vs. mating: Z = 11.83, p < 0.0001). Mating males skewed larger than the total population of foraging males (Mann-Whitney test: U = 1439, p < 0.0001). There was no difference in the mean head widths of digging and fighting males (unpaired t-test: t = 1.55, df = 135, p = 0.12).

Patrolling and foraging male head widths increased over the course of the morning (Supplementary Figure 2; linear regression, patrolling: [head width] = 0.095 [time of day] + 4.19, F = 23.65, df = 408, R^2^ = 0.05, p < 0.0001; foraging: [head width] = 0.097 [time of day] + 4.08, F = 7.90, df = 201, R^2^ = 0.03, p < 0.0001); hovering and mating male head widths were constant across the morning (hovering: F = 1.57, df = 88, p = 0.21; mating: F = 0.43, df = 57, p = 0.51).

### Historical comparison of mean male head widths from 1974 to 2022

Reported head widths for males foraging, hovering, patrolling, or mating at the Blue Point Bridge site from 1974 to 2022 can be found in Table 1, with associated references for the historical data.

**Table 1.**
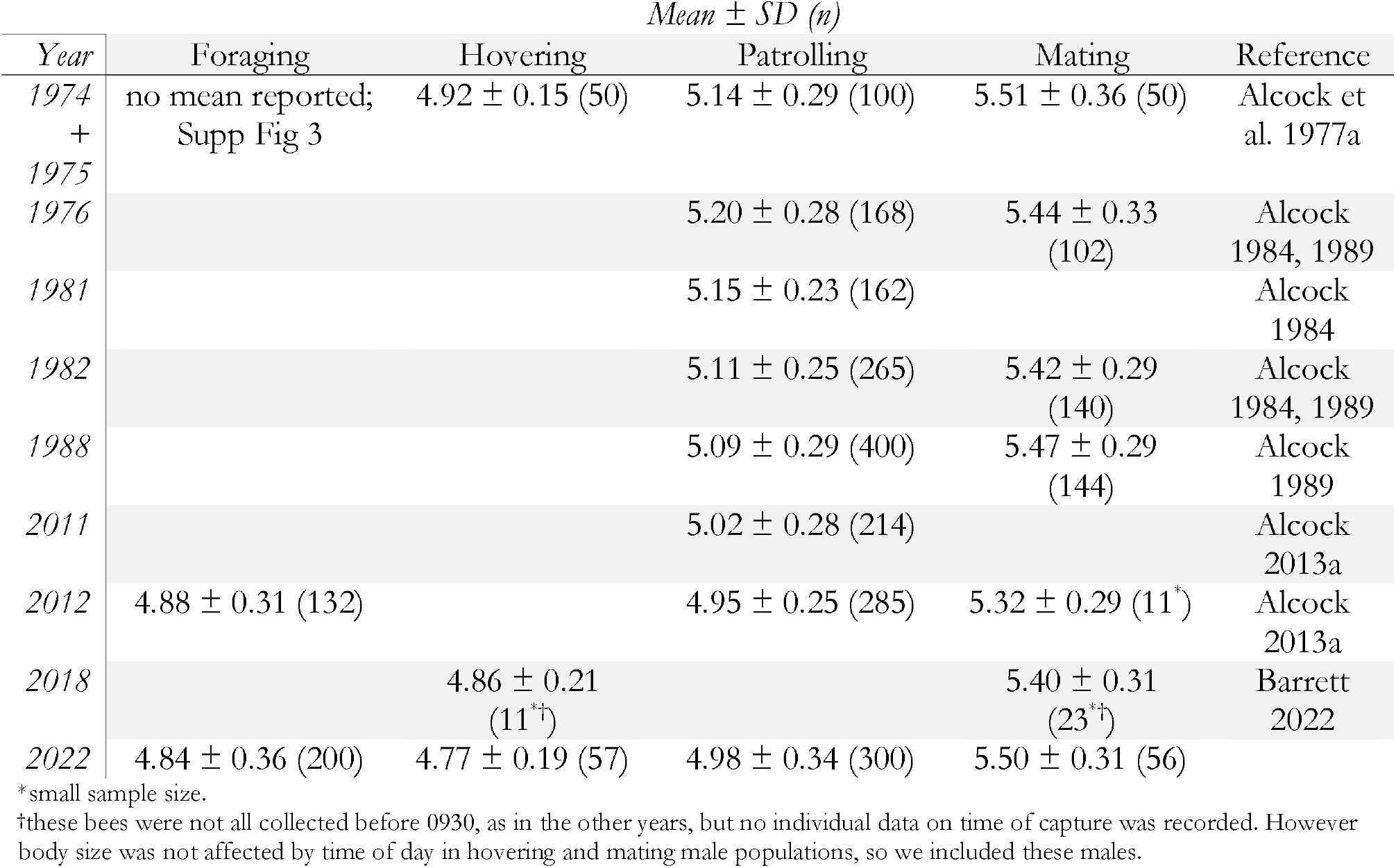
Reported means ± SD of males at Blue Point Bridge before 930 from 1974 – 2022.

The mean head width of the foraging males (n = 100; representing the total population of males), was not reported in Alcock et al. 1977a. However, comparing the distribution of head widths from 1974/1975 to our 2022 data suggests significant declines in overall male head width (Figure 2B & 2C). The most frequent head width class declines from 5.0 – 5.22 mm to 4.6 – 4.8 mm (which was previously the smallest size class). The smallest size class observed in the male population decline from 4.6 – 4.8 mm to 4.0 – 4.2 mm. The two largest size classes (5.8 – 6.2 mm) were lost entirely from the foraging male distribution despite surveying twice the number of foraging males in 2022 compared to 1974/1975. These two size classes decreased from ∼30% of all mating events in 1974/1975 to ∼10% in 2022, demonstrating a significant decrease in the largest size large-morph males.

**Figure 2.**
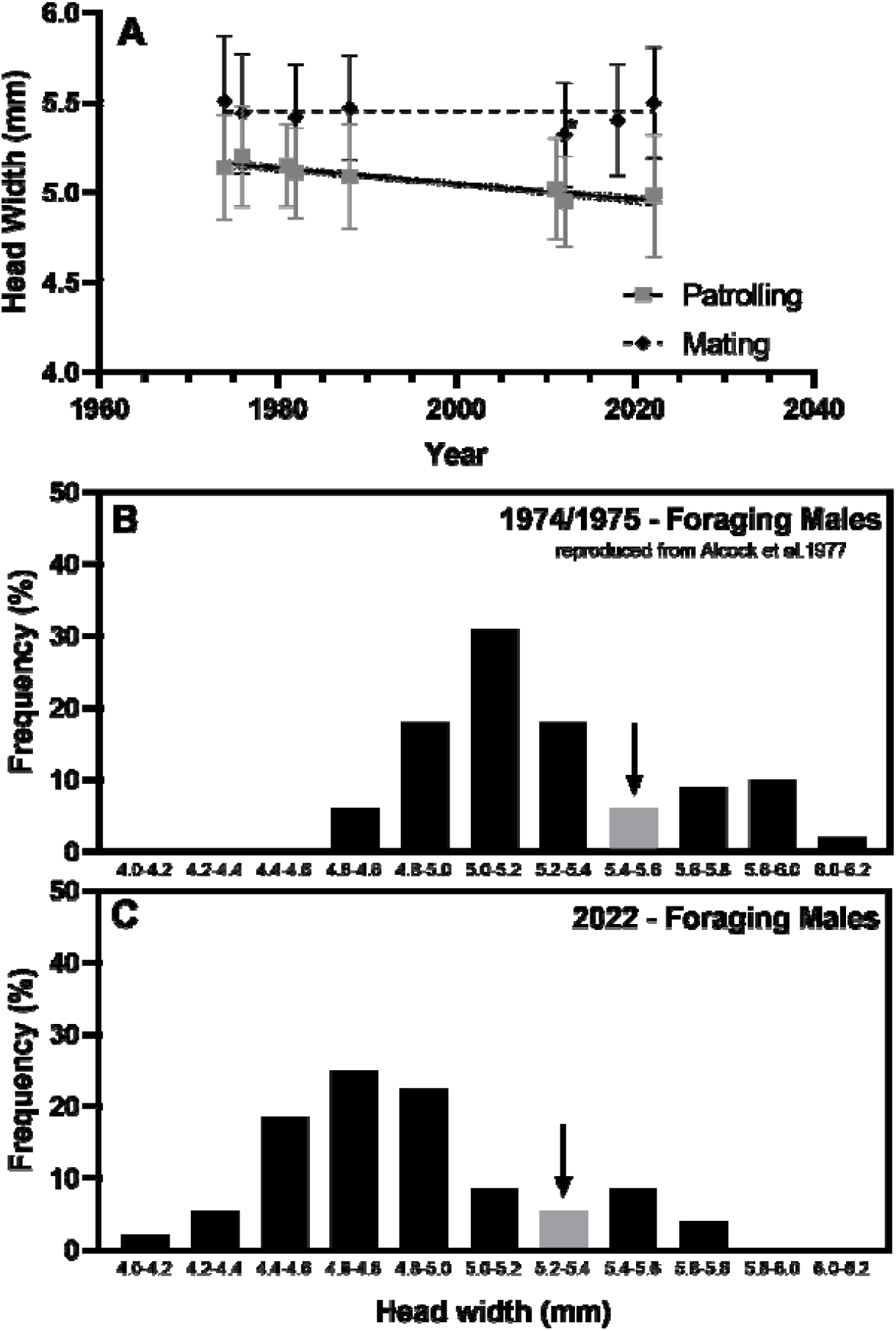
Changes in head widths of patrolling, foraging, and mating males from 1974 to 2022. A) Patrolling male head widths (gray dots, solid line, n = 300) decreased over time at the Blue Point Bridge site from 1974 to 2022 (Figure 2; linear regression, [head width] = -0.004 [year] + 13.74, F = 131.2, df = 1892, R^2^ = 0.06, p < 0.0001). Mating male head widths (black diamonds, dashed line to demonstrate non-significance, n = 56) did not change from 1974 to 2022 (F = 0.36, df = 501, p = 0.55). Black dotted lines = 95% confidence interval. *note: small sample size (n = 11) for this year (Alcock 2013a). B) Distribution of head widths of foraging males (n = 100) reported in Alcock et al. 1977a shows larger median head widths than C) the distribution of foraging males (n = 200) collected in 2022. All males collected before 930 am. The arrow represents the rare intermediate males (5-6%; gray column, local minimum); to the left of this head width class are the small-morph males, to the right are the large-morph males.

Additionally, there were within-morph body size declines for both morphs. The most frequent head width for small-morph males (to the left of the arrow in Figure 2B/2C) in 1974/1975 was 5.0 – 5.2 mm; by 2022, the peak was 4.6 – 4.8 mm (previously, the smallest size category for all males). For large-morph males (to the right of the arrow) the peak frequency also decreased, from 5.8 – 6.0 mm to 5.4 – 5.6 mm.

Behaviorally-linked data corroborate this decrease in body size. Patrolling male head widths, which represent males from both morphs (but is skewed towards larger small-morph, and large-morph, males compared to the overall population of foraging males), decreased by 3.72% at the Blue Point Bridge site from 1974 to 2022 (Figure 2A, Supplementary Figure 3B; linear regression, [head width] = -0.004344 [year] + 13.74, F = 131.2, df = 1892, R^2^ = 0.06, p < 0.0001). There was a 3.05% decrease in mean hovering male head width from 1974/1975 to 2022 (Supplementary Figure 3A, unpaired t-test; t = 4.49, df = 105, p < 0.0001); hovering is exclusively performed by small-morph males, corroborating the decrease in head width shown in the foraging male frequency distribution data.

Small-morph males increased from 73% of the total male (foraging) population in 1974/1975 to 82% in 2022. Large-morph males (to the right of the arrow) decreased from 21% to 13%; intermediates were rare and represented only 5-6% of all males in each year.

## Discussion

The mean head width of *C. pallida* males engaged in foraging (e.g., total population), hovering, and patrolling has declined at Blue Point Bridge over the past five decades. The most frequent head width class among all males has shifted from 5.0 – 5.2 mm in 1974 to 4.6 – 4.8 mm in 2022 (foraging males, Figure 2B,C). Within-morph, both large- and small-morph males have seen a similar decline of 0.4 mm in their most frequent head width.

Time-of-day affects body size for patrolling and foraging male populations, but not mating or hovering males – foraging and patrolling males were smaller early in the morning. This may relate to the thermal adaptations of the males: the darker coloration of small-morph males may allow them to heat up their flight muscles faster in the cool early mornings, while the lighter coloration of large-morph males may keep them cooler later in the morning (Barrett et al. 2022, Barrett & O’Donnell 2022). A similar coloration effect, which keeps the flight muscles at an optimal temperature while balancing convective cooling and shortwave radiative heat gain, is seen in male butterflies with different mating behaviors (Van Dyck & Matthysen 1998). Alternately, small-morph males may simply emerge earlier in the day to avoid displacement by large-morph males in fights, as Alcock (2009) hypothesized for size-dimorphic *Amegilla dawsoni* (Hymenoptera: Apidae) males that use similar ARTs (but do not differ in coloration between the morphs). However, our data suggest this is unlikely in *C. pallida* – males mating earlier in the morning are not any smaller in body size, suggesting that small-morph males are not avoiding displacement by larger males simply by ‘getting up’ early. Therefore, rather than sexually-selected early emergence, we favor the physiological limitations on flight muscle temperature hypothesis. Future studies may also look for temporal variation in male body size across the emergence season (Alcock 2009).

We also found that small-morph males were a larger proportion of the overall population in 2022 compared to 1974/1975 (from 73% to 82% of the total foraging population). Large-morph males appear to be slowly declining within the overall population, representing a potential threat to the longevity of the ART system in this species. Despite their numerical decline, the large-morph fitness advantage continues to hold: head widths of the mating male population have remained constant over time, despite a significant decline in the frequency of males of this larger size class in the population overall (Alcock et al. 1977a, 1977b, Alcock 1984, Alcock 1995, Alcock 2013a). However, the largest large-morph male size class (5.8 – 6.2 mm) decreased from nearly 30% of mating events to only 10% in 2022, providing further evidence that the large-morph males are declining. Changes in body size over time have been reported for female bees in temperate regions (Bartomeus et al. 2013, Scheper et al. 2014, Oliveira et al. 2016, Nooten & Rehan 2020). Unlike females, males were not reported to decline in body size in the Netherlands; Oliveira et al. (2016) propose that fitness advantages associated with larger male body size may prevent similar declines in males. However, the fitness advantages of larger male body sizes are unclear or nonexistent in many bee species, particularly those without male-male competition (Alcock 2013b). Thus, in combination with our results on *C. pallida* (where large-morph males are declining in frequency), this seems like an unlikely explanation for the trends observed across bees so far. Further studies in other systems with large-male fitness advantages (Kukuk 1996, Danforth 1991, Alcock 1983, Alcock 1994, Alcock 1997, Paxton 2005) would be beneficial to test this hypothesis.

Sex-biased resource allocation is common in many Hymenoptera (e.g., O’Neill & O’Neill 2009), generating male bees that are typically smaller than the females of their species (Shreeves & Field 2008). This may help explain the differences observed in male body size trends between *C. pallida* (where this is not the case) and the other studied species in Oliveira et al. (2016). Low quantity or quality larval nutrition significantly reduces survival and impacts adult physiology (Lawson et al. 2020, Nicholls et al. 2021), resulting in reduced mating and foraging success (Muller et al. 2015, Xie et al. 2015). The negative fitness consequences of small body sizes may cause a species-specific lower bound on resource allocation per offspring. As male bees are already closer to that species-specific lower bound in resource allocation, this may prevent male bees of most species from declining as much in body size relative to females. *C. pallida* is unique in that males can be the same size or larger than females, allowing for more significant size declines in the male population. This phenomenon may also explain the shift in the proportion of large-versus small-morph males, as there is more room in the *C. pallida* system to decrease the high end of the male body size spectrum. Alternately, other life history or ecosystem characteristics between the bees studied in Oliveira et al. (2016) and *C. pallida* may be responsible for variation in reported male body size declines. As a large-bodied, solitary, desert-adapted bee species with narrow phenological and dietary breadth, *C. pallida* may be particularly susceptible to human-activity induced declines. Hypotheses for bee body size declines include: 1) habitat simplification or agricultural intensification, 2) climate-induced phenological mismatches between bees and host plants, or changes in total resource availability (e.g., lower floral abundance due to more persistent droughts), and 3) increasing temperatures during development (Chole et al. 2019; Sánchez-Bayo and Wyckhuys 2019). Although we do not test these hypotheses in this report, we provide contextual information that may support particular causes of *C. pallida* body size declines.

The area where *C. pallida* have been studied is surrounded by land managed by the United States Forest Service, with minimal development since the 1970s. It is unlikely that habitat simplification or agricultural intensification is responsible for body size declines. Soil significantly buffers ground temperature fluctuations at 10 cm depth (Parton & Logan 1981, Cane & Neff 2011; approximately *C. pallida* nest depth, Alcock et al. 1976a), which should lessen the impact of increasing temperatures on development and thus body size.

Resource limitation may occur via a combination of phenological mismatch and climate-induced reductions in floral availability. Phenological mismatch, which may arise more frequently for ground-nesting bees with short foraging periods (Stemkovski et al. 2020), is a possible driver of reduced body sizes in *C. pallida*. Though inconsistently documented, it is likely that the emergence dates of *C. pallida* have shifted earlier since the 1970-80s. The female flight season was documented as late May to mid-June in Alcock et al. (1976a); this year (2022), the beginning of the flight season was late April.

More data are available on dates of mating aggregations than female foraging behavior at this site and support the idea of a two week advance in phenology since the 1980s. In 1982, peak emergence was between April 30 – May 14 (Alcock 1984). Peak emergence is even earlier in 2012: Alcock (2013a) suggests peak activity around April 25-May 4. By 2022, we found peak mating between April 20-May 4, an additional five days sooner. Generalist bees in the Northeastern United States have advanced in phenology by ∼10 days over the last 130 years, with most of that advance occurring after 1970 (Bartomeus et al. 2011); this aligns with shifts observed in the *C. pallida* mating aggregation activity. However, phenological mismatch is unlikely to be the sole cause of body size declines – studies of native bees have demonstrated that they largely advance at a pace similar to their host plants (Bartomeus et al. 2011). In addition, data on the flowering periods of the three most common host plants for *C. pallida*, collected by the Arizona-Sonora Desert Museum in Tucson from 1983-2009, does not suggest an obvious mismatch in flowering and emergence dates (Arizona-Sonora Desert Museum, 2022).

Resource limitation could affect male body size by 1) decreasing female body sizes (maternal and offspring body sizes are linked in *C. pallida;* Alcock 1979) or 2) altering female resource allocation decisions. Unfortunately, no data are available on female *C. pallida* body sizes at Blue Point Bridge prior to 2022, and this relationship is expected to be population-dependent (Alcock 1979). However, body size can affect female flight velocity and pollen carrying capacity (Müller et al. 2006, Everaars et al. 2018), which may have dramatic effects on the number or size of offspring of different types in *C. pallida*. Alcock et al. (1977a; Alcock 1979) propose that a females’ assessment of the likelihood of nest failure due to parasitism or resource limitation may also affect her decision to allocate for more, smaller (risk-averse strategy) versus fewer, larger male offspring. Notably, resource limitation seems more likely than parasitism: rates of parasitism by bee flies, blister beetles, and mutillids are notably low (Alcock 1979), and no kleptoparasite bees are known to parasitize *C. pallida* nests (Rozen & Buchmann 1990).

Irrespective of the mechanism responsible for declines in overall male *C. pallida* body size, reductions in the frequency of large-morph males, specifically, over time may have many important consequences for the species. First, the opportunity to pass on genetic information may become increasingly limited to an ever-smaller pool of males capable of winning competitions (the largest males may mate multiple times; Alcock 1995). However, given that small-morph males will patrol and mate with females whenever large-morph males are not around to compete with them, this outcome may be unlikely in the *C. pallida* mating system.

Second, the stability of the *C. pallida* male ART system, which likely relies on competing selective forces related to nesting density, female provisioning behavior, and male mating success (Alcock 1979), may be threatened by declines in the species’s mean body size and the decreasing frequency of large-morph males. The impacts of human activities often consider biodiversity loss at the level of the species, however losses in intraspecific diversity (variation in behavior or morphology) are currently underexplored (see Bolnick et al. 2003). ARTs represent the functional potential a species has to adapt to a changing environment (Oliviera et al. 2016); losses in ecologically functional intraspecific diversity, which might occur if size-based ARTs are destabilized, should be considered when evaluating the impact of human activities. Additional work following *C. pallida* male and female body sizes at the Blue Point Bridge population may allow for the mechanisms of body size decline in bees broadly, and *C. pallida* specifically, to be further tested, as human-induced climate and landscape modifications continue to drive morphological and behavioral changes in a variety of species.

## Acknowledgements

We thank John Alcock for his record of *C. pallida* head widths at Blue Point over the years, and Stephen Buchmann for his kindness and valuable *C. pallida* wisdom, which have made *C. pallida* an accessible system for many early career bee researchers; his comments also greatly improved an early draft of this paper. We thank Victoria Morgan for helping collect head widths on May 8^th^, 2022. We also thank the editor, and two anonymous reviewers, for thoughtful comments that improved this manuscript significantly. Support was provided to MGJ by Friends of the Sonoran Desert and MB by the National Science Foundation. MB is an NSF postdoctoral fellow (2109399): any opinions, findings, conclusions, or recommendations expressed in this manuscript are the authors, and do not necessarily reflect the views of the NSF.

## Supplemental Figures

**Supplementary Figure 1.**
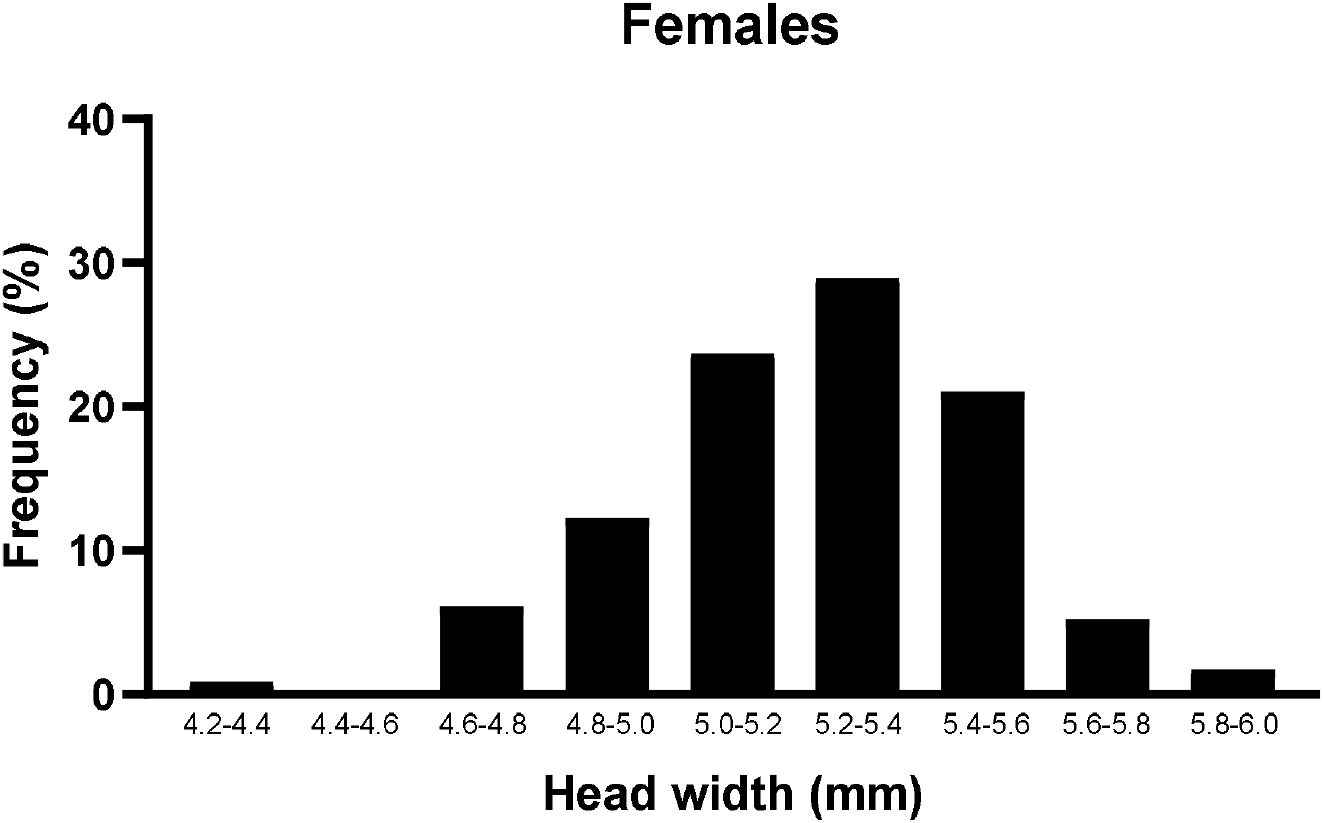
Distribution of female head widths in 2022. Distribution of head widths of females (n = 114) collected while mating, foraging, and nesting from before 1130 on April 20 – May 8, 2022, at Blue Point Bridge, Arizona.

**Supplementary Figure 2.**
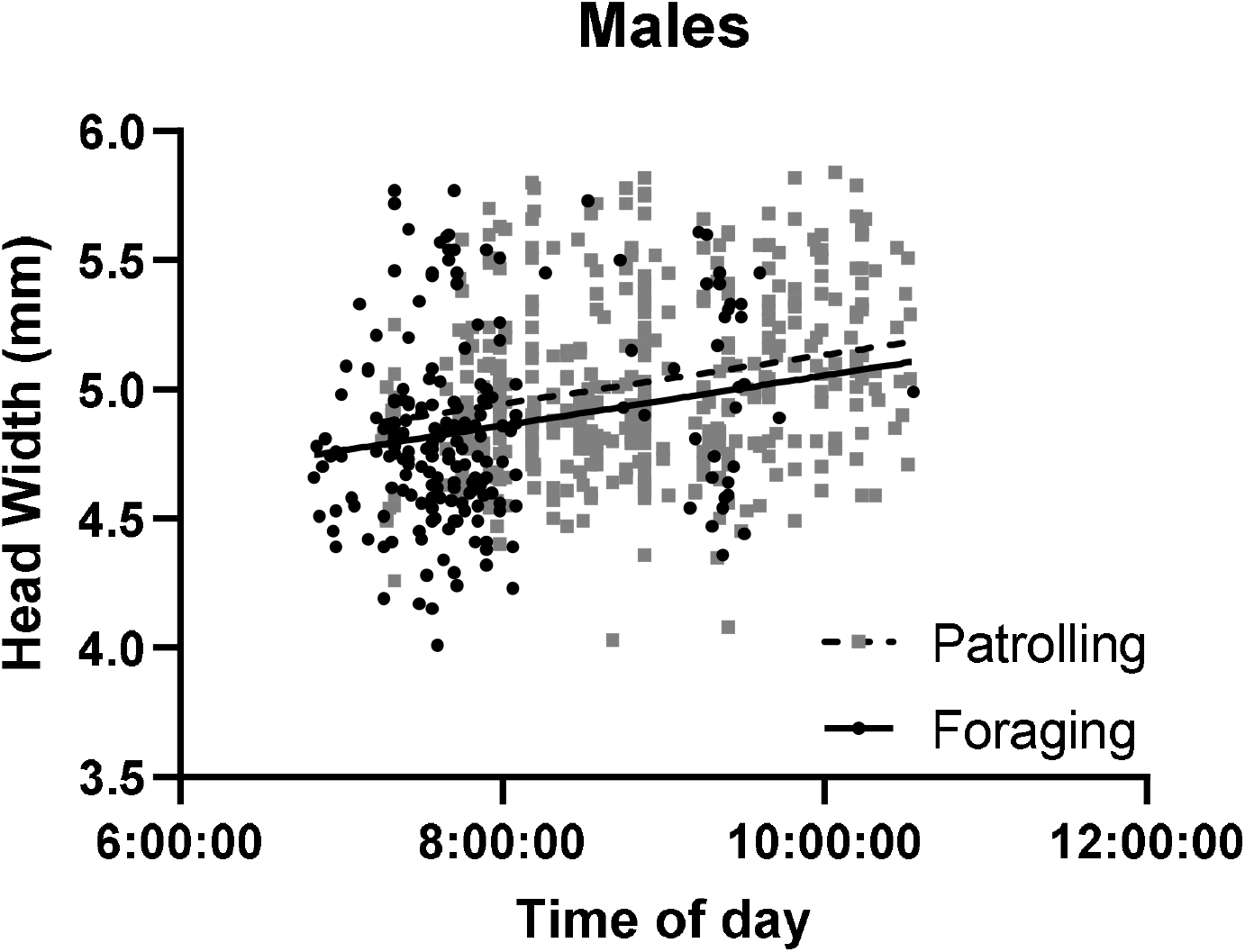
Increase in patrolling or foraging male head widths from 0700 to 1130. Head width in the populations of males patrolling (gray squares, dashed line, n = 410) and foraging (black dots, solid line, n = 203) increased over the course of the morning (Supplementary Figure 2; linear regression, patrolling: [head width] = 0.095 [time of day] + 4.19, F = 23.65, df = 408, R^2^ = 0.05, p < 0.0001; foraging: [head width] = 0.097 [time of day] + 4.08, F = 7.90, df = 201, R^2^ = 0.03, p < 0.0001).

**Supplementary Figure 3.**
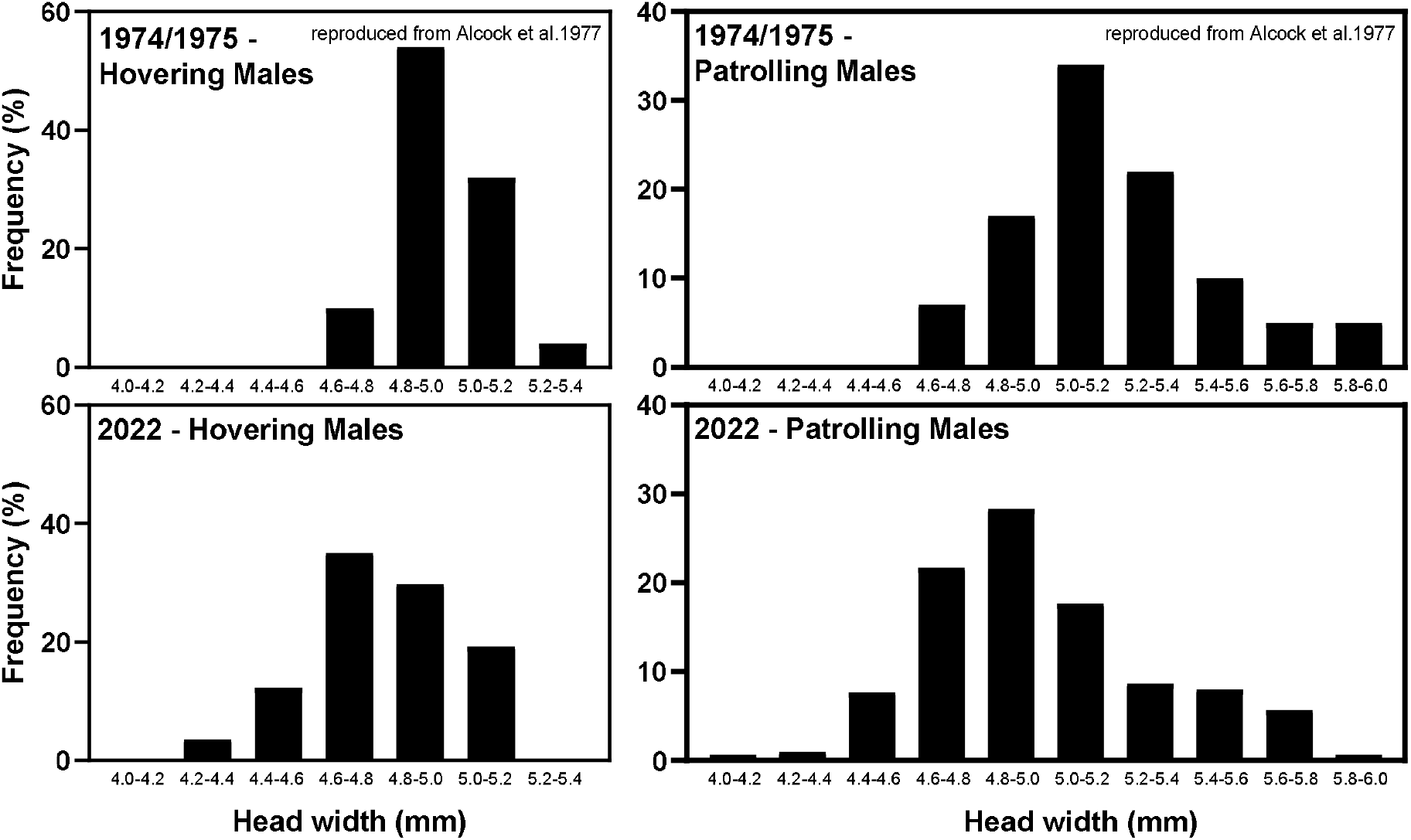
Changes in head width distributions of hovering and patrolling males from 1974 to 2022. Head width distributions for hovering (n = 57) and patrolling (n = 300) males at Blue Point Bridge are skewed towards smaller body sizes in 2022 as compared to 1974/1975 (n = 50 hoverers, 100 patrollers; data reproduced from Alcock et al. 1977a). All males collected before 930 am.

**Supplementary Figure 4.**
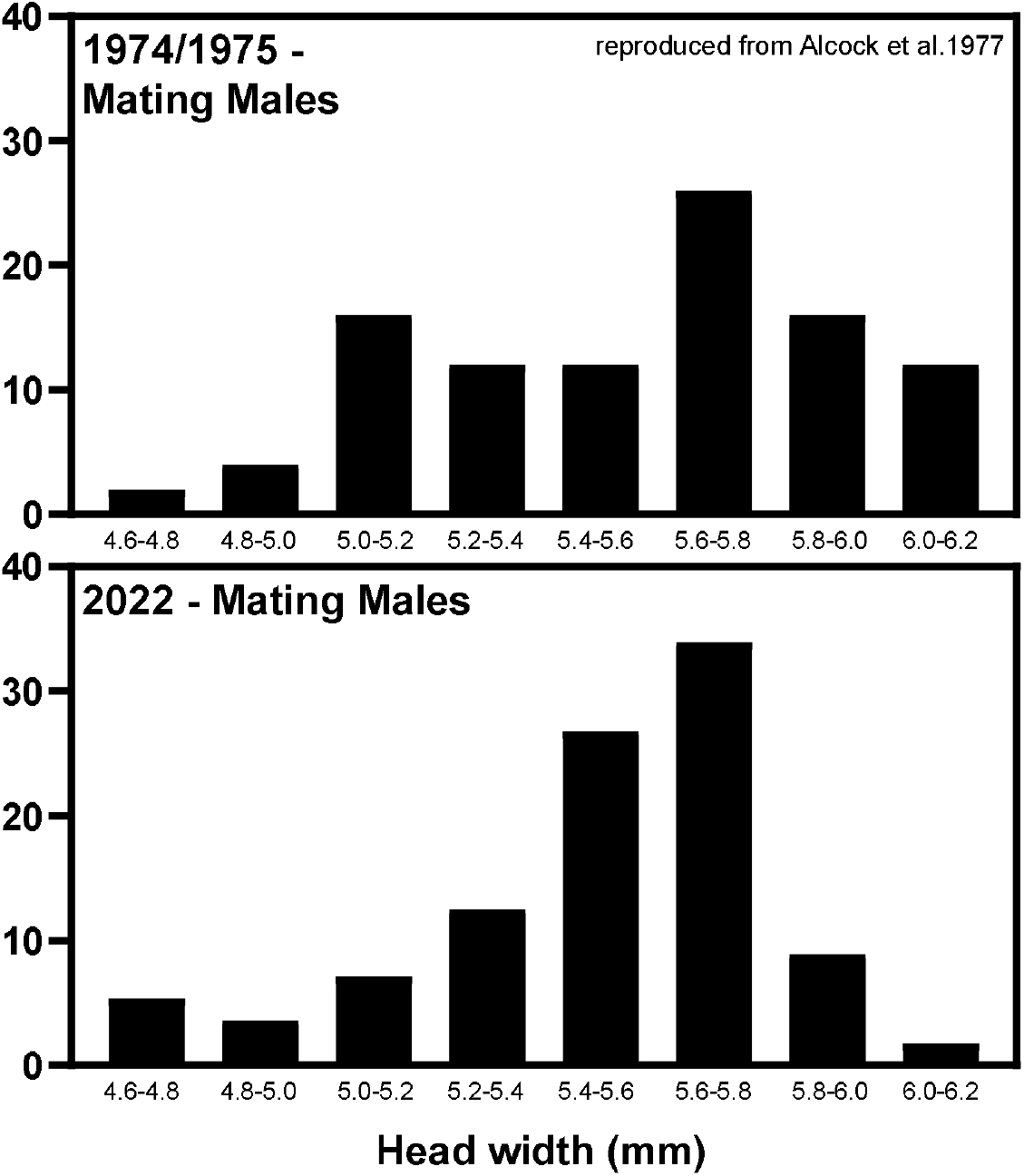
Similar head width distributions of mating males from 1974 to 2022. Head width distributions for mating males at Blue Point Bridge demonstrates the population of males that obtain mating opportunities have relatively similar body sizes in 2022 (n = 56) as compared to 1974/1975 (n = 50; data reproduced from Alcock et al. 1977a). All males collected before 0930.

**Supplementary Table 1.**
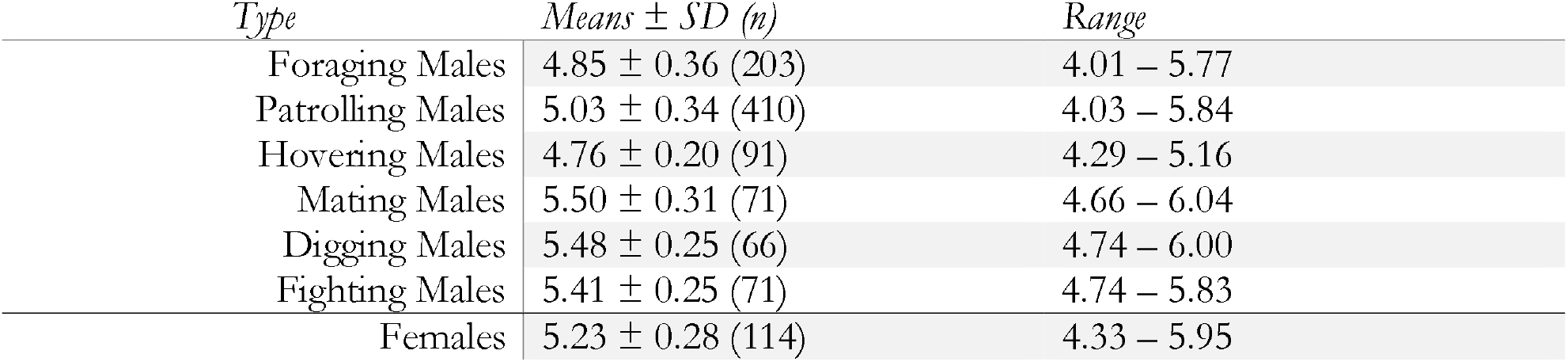
Head widths of *C. pallida* collected in April - May 2022 at Blue Point Bridge before 1130 am (n = 921 male, 114 female).

| Type | Means ± SD ($n$) | Range |
| --- | --- | --- |
| Foraging Males | 4.85 ± 0.36 (203) | 4.01 – 5.77 |
| Patrolling Males | 5.03 ± 0.34 (410) | 4.03 – 5.84 |
| Hovering Males | 4.76 ± 0.20 (91) | 4.29 – 5.16 |
| Mating Males | 5.50 ± 0.31 (71) | 4.66 – 6.04 |
| Digging Males | 5.48 ± 0.25 (66) | 4.74 – 6.00 |
| Fighting Males | 5.41 ± 0.25 (71) | 4.74 – 5.83 |
| Females | 5.23 ± 0.28 (114) | 4.33 – 5.95 |

